# Inhibition of translation termination by Drosocin, an antimicrobial peptide from fruit flies

**DOI:** 10.1101/2022.12.11.519952

**Authors:** Kyle Mangano, Dorota Klepacki, Irueosa Ohanmu, Chetana Baliga, Weiping Huang, Alexandra Brakel, Andor Krizsan, Yury S. Polikanov, Ralf Hoffmann, Nora Vázquez-Laslop, Alexander S. Mankin

## Abstract

A 19-amino acid long <u>p</u>roline-<u>r</u>ich <u>a</u>nti<u>m</u>icrobial <u>p</u>eptide (PrAMP) Drosocin (Dro) is encoded in the fruit fly genome. Native Dro is glycosylated at a specific threonine residue, but the non-glycosylated peptide retains antibacterial activity. Dro shows sequence similarity to several other PrAMPs that bind in the ribosomal nascent peptide exit tunnel and inhibit protein synthesis by varying mechanisms. However, the target and mechanism of action of Dro remain unknown. Here we show that the primary mode of Dro action is inhibition of termination of protein synthesis. Our in vitro and in vivo experiments demonstrate that Dro stalls ribosomes at stop codons, likely sequestering class 1 release factors associated with the terminating ribosome. As the result, Dro strongly promotes readthrough of stop codons at subinhibitory concentrations. The elucidated mode of Dro action allows assigning it as the second member of the type II PrAMPs, of which only one representative, the antimicrobial peptide apidaecin (Api) produced by honeybees, was previously known. However, despite its functional similarity with Api, Dro interacts with the target in a markedly distinct way. The analysis of a comprehensive single-amino acid substitution library of endogenously expressed Dro variants shows that binding to the ribosome involves interactions of multiple amino acid residues distributed through the entire length of the PrAMP. Our data further show that the ribosome-targeting activity of non-glycosylated Dro can be significantly enhanced by single amino acid substitutions illuminating directions for improving its antibacterial properties.

## Introduction

Antimicrobial peptides (AMPs) are important components of the innate immune system of higher organisms helping them to suppress infections caused by bacterial pathogens ^1^. The best-studied AMPs kill bacteria by attaching to the bacterial cytoplasmic membrane, disrupting its integrity and causing cell lysis ^2^. However, some AMPs penetrate the bacterial cell and inhibit its growth by acting upon intracellular targets ^3^. Such non-lytic AMPs are particularly interesting because besides inspiring the development of clinically useful antibiotics, they can serve as probes for elucidating operation of cellular mechanisms they act upon. Proline-rich AMPs (PrAMPs) belong to the latter class ^4-7^.

The known PrAMPs, produced by insects and mammals, range in size from ∼15 to ∼60 amino acids ^8,9^. However, their pharmacophore (the sequence critical for the on-target activity) is usually limited to a smaller segment; in some cases peptides with as few as 12 amino acid residues may retain activity ^10,11^. Most of the known PrAMPs are produced as unmodified peptides, but few carry posttranslational modifications ^12,13^.

While initially it was proposed that PrAMPs stop cell growth by inhibiting the activity of the protein chaperone DnaK ^14-16^, subsequent discovery that DnaK-lacking mutants remain sensitive to PrAMPs ^17-19^ inspired further investigation of the mode of action of these antibiotics. More recent biochemical and structural studies showed that PrAMPs could bind to the ribosome and inhibit its functions ^19-25^. The eventual demonstration that ribosomal mutations confer resistance to PrAMPs firmly established the ribosome as the primary target of PrAMPs in bacteria ^11,23,25^.

Specific peptide transporters import PrAMPs into the cytoplasm of the bacterial cell ^26,27^. There, they bind to the ribosome. PrAMP’s binding site is located in the in the large ribosomal subunit (50S in bacteria), specifically, in nascent peptide exit tunnel (NPET) ^21-25^, which serves as the passageway for the growing polypeptide assembled in the peptidyl transferase center (PTC). The PTC is also responsible for the release of the completed protein from the P-site tRNA in the reaction involving class-1 release factors (RF1 and RF2 in bacteria) that bind to the ribosome at the stop codons. According to their binding mode and mechanism of action, the examined PrAMPs fall into two major categories ^9^. Most of them, including oncocin, bactenecin, pyrrhocoricin and metalnikowin, belong to type I. The C-termini of type I PrAMPs protrudes down the NPET while their N-termini invade the A site of the PTC catalytic center ^21-24^ (**Fig. 1a**). The PrAMP-associated ribosome can form the initiation complex at the mRNA start codon but cannot accommodate the first elongator aminoacyl-tRNA and catalyze formation of the first peptide bond. As a result, type I PrAMPs stop protein synthesis at translation initiation stage by arresting ribosomes at mRNA start codons ^21-24,28,29^. The only known PrAMP belonging to type II, apidaecin (Api) ^4^, acts in a drastically different way. The synthetic apidaecin derivative Api137 ^30^ also binds in the NPET, but in contrast to type I PrAMPs, in an orientation that mimics that of the nascent polypeptide, i.e., with the N-terminus stretched down the tunnel and the C-terminus bordering the PTC ^25,31^ (**Fig. 1a**). Api acts upon the ribosome that has reached the stop codon and is still associated with RF1 or RF2 ^25,32^. Once the newly made polypeptide vacates the ribosome, Api diffuses up the NPET and its penultimate arginine residue (R17) bonds with RF. In addition, Api’s C-terminal leucine (L18) interacts with the 3’ ribose of the P-site-bound deacylated tRNA ^25^. As a result, Api freezes the ribosome at the stop codon, and sequesters class-1 RFs ^25,31^. Since in the bacterial cell there is an excess of ribosomes over RFs, the free RF pool is rapidly depleted and when the rest of the ribosomes reach the stop codons, they are unable to release the completed proteins. This leads to ribosome stalling in a pre-release state and eventual stop codon bypass by aminoacyl-tRNA misincorporation or frameshifting ^11,25,32^.

**Fig. 1:**
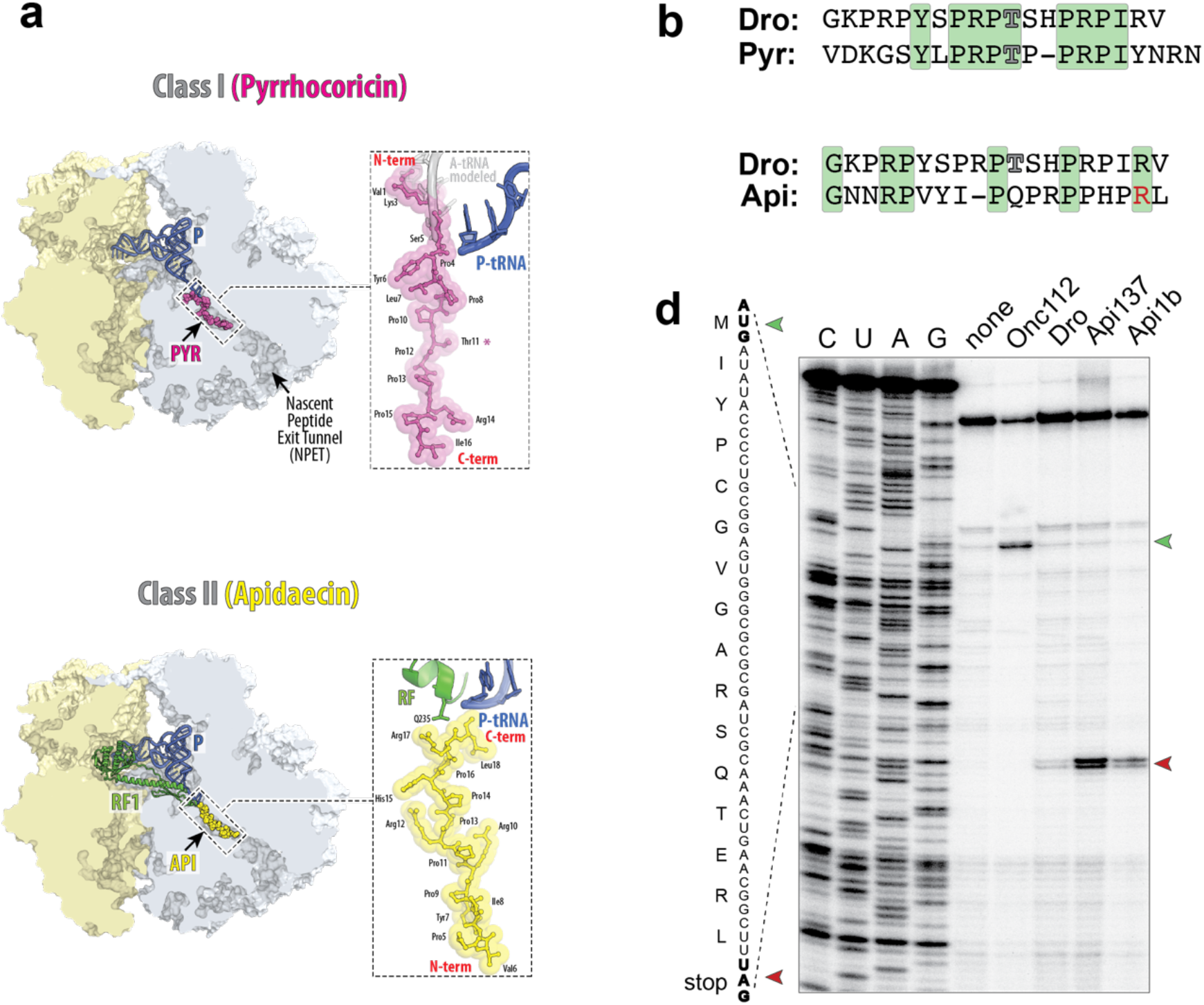
The PrAMP Drosocin stalls ribosomes at stop codons during in vitro translation. **a**, Comparison of binding of type I PrAMP pyrrhocoricin (Pyr) (PDB 5HD1 ^23^) and type II PrAMP apidaecin (Api) (PDB 5O2R ^25^) to the bacterial ribosome. **b**, The Drosocin (Dro) amino acid sequence, aligned with that of type I PrAMP Pyrrhocoricin (Pyr) or type II PrAMP Apidaecin (Api). Residues identical between the compared PrAMPs are highlighted in green. The glycosylated Thr (T) residues of Dro and Pyr are indicated by outlined characters. The functionally critical penultimate Arg (R) residue of Api is highlighted in red. **c**, Toeprinting analysis of the ribosome arrest at the UAG stop codon of a model ORF derived from the *E. coli yrbA* gene due to the addition of synthetic PrAMPs Onc112, Dro, Api137, or Api1b. The control reaction where no PrAMPs were added is labeled as “none”. The toeprint bands corresponding to ribosomes arrested at the start or stop codons of the ORF are marked with green or red arrowheads, respectively. Sequencing reactions are labeled as C, U, A, G.

None of the other characterized PrAMPs are known to act in an Api-like fashion. However, it was noted that drosocin (Dro), a PrAMP encoded in the *Drosophila melanogaster* genome, shows some sequence similarity to Api and may compete with Api for binding to the ribosome ^20^. In the 19-amino acid long sequence of Dro, GKPRPYSPRP<u>T</u>SHPRPIRV, Thr11 (underlined) is O-glycosylated with N-acetylgalactosamine and galactose ^12^. The glycosylation of Thr11 stimulates the antibacterial activity of Dro, reducing its MIC by up to 8-fold compared with the unmodified peptide ^12,33,34^, but it remains unclear whether this posttranslational modification is important for the uptake of the PrAMP or its interaction with the intracellular target. Dro shows promising activity against a range of Gram-negative and some Gram-positive bacteria, including clinically relevant species ^33-37^.

While Dro is one of the first discovered PrAMPs, little is known about its mode of action. Although Dro binds to the ribosome with a moderate affinity ^20^, it remains unclear whether it inhibits cell growth by interfering with translation. Furthermore, even if the ribosome is indeed the primary target of Dro, it is unclear which step of translation might be affected and through which mechanism.

Here we unraveled the mechanism of action of Dro in the bacterial cell and identified the amino acid residues critical for target engagement and activity. The results of our biochemical and genetic experiments established Dro as an efficient inhibitor of translation termination. We further show that the unmodified Dro retains its inhibitory activity when expressed directly in the bacterial cell. By exploiting endogenously expressed Dro, we surveyed the entire sequence space of single amino acid substitutions and identified the amino acid residues critical for the ribosome-targeting activity of this PrAMP. Furthermore, we discovered mutant variants which exhibit superior activity in comparison to the unmodified wild-type (wt) Dro. Our findings establish that a range of short peptides with diverse sequences can interfere with the termination stage of translation by binding in the ribosomal NPET.

## Results

### Dro arrests the ribosome at the stop codon of the ORF

The presence of the glycosylated threonine in naturally produced Dro and in the type I PrAMP pyrrhocoricin ^13^, with which Dro shows moderate sequence similarity (**Fig. 1b**), may suggest that, like type I PrAMPs, Dro could invade with its N-terminal segment the PTC A site and inhibit formation of the first peptide bond ^21,23^ (**Fig. 1a**). On the other hand, the comparable sequence similarity between Dro and Api (**Fig. 1b**) leaves it possible that Dro’s mode of binding and inhibition could be similar to that of Api ^20^ (**Fig. 1a**). To establish the genuine mechanism of Dro action, we tested its effects on protein synthesis in a cell-free translation system. Because glycosylation of the Thr11 residue is neither strictly required for the antimicrobial activity of Dro ^12,33,34^ nor is critical for the interaction of the PrAMP pyrrhocoricin with the ribosome ^23^, we avoided the rather cumbersome synthesis of the glycosylated peptide and carried out all our experiments with the unmodified synthetic Dro.

In agreement with published results ^20,38^, in vitro expression of the GFP reporter is significantly inhibited by type I PrAMP Onc112, which interferes with initiation. In contrast, type II PrAMP Api137, which affects termination and allows at least one round of mRNA translation, inhibits GFP expression only moderately, even at high concentrations (**Extended Data Fig. 1a**). Like Api137, Dro inhibited GFP expression rather inefficiently. In agreement with the known mechanism of action, the addition of RF1 exacerbated inhibition of translation by Api137, but not by Onc112. While added RF1 had only mild effect on Dro-mediated inhibition of translation at lower concentrations of the PrMAP, at higher concentration, Dro diminished GFP expression more efficiently in the RF1-augmented reaction indicating that Dro’s action might be related to termination of translation (**Extended Data Fig. 1a**).

To further explore the possibility that Dro interferes with translation termination, we tested its effects on the progression of ribosomes through mRNA by toeprinting, a technique that maps inhibitor-induced ribosome stalling sites during in vitro translation ^39,40^. Consistent with previous reports ^21,23^, type I PrAMP Onc112 arrested the ribosomes at the start codon of the open reading frame (ORF) whereas Api137, or its native prototype Api1b, caused translation arrest at the mRNA’s stop codon (**Fig. 1c**) ^25^. Addition of Dro had no effect on translation initiation, but a distinct toeprinting band corresponding to the ribosome arrested at the UAG stop codon appeared (**Fig. 1c**). Thus, consistent with the results of in vitro GFP translation (**Extended Data Fig. 1a**), toeprinting analysis further supports our conclusion that Dro inhibits termination of translation.

Replacement of the UAG stop codon with UGA significantly reduced Api-dependent translation arrest and diminished Dro-induced ribosome stalling (**Extended Data Fig. 1b**). The RF2 present in the toeprinting reaction originates from the *E. coli* K12 strain and carries the endemic Ala246Thr mutation ^41^ which, as we showed previously, renders it resistant to the action of Api ^25^. A weaker response of the K12 variant of RF2 to Dro argues that the Ala246Thr substitution also diminishes sensitivity to this PrAMPs.

### Dro inhibits bacterial growth by interfering with translation termination

Although our in vitro experiments show that Dro stalls ribosomes at stop codons (**Fig. 1**; **Extended Data Fig. 1**), these results do not prove that inhibition of translation termination is the primary mode of Dro action against bacterial cells, especially considering that Dro was previously shown to interfere with the activity of the DnaK chaperone ^14,42^. To identify the target of Dro action in bacteria, we tested its effect upon several mutants. We first compared Dro’s minimal inhibitory concentrations (MICs) against the *E. coli* K12 strain BW25113 and its Δ*dnaK* mutant. Although parental BW25113 strain is fairly tolerant to Dro due in part to the intrinsic Ala246Thr mutation in the RF2 gene ^41^, it can be readily inhibited with 32-64 μg/mL of Dro. Deletion of the *dnaK* gene did not alter the Dro MIC, thereby excluding DnaK as the main target. We further determined Dro’s MIC against the mutants of the *E. coli* B strain BL21 carrying alterations in ribosomal protein uL16 (the R18C substitution) or in RF2 (the R262C or Q280L mutations), isolated previously for their resistance to Api ^25^. (Of note, RF2 of the wt BL21 cells has an ‘unmutated’ Ala246 conserved in other enterobacteria). All the three tested mutants showed significantly increased resistance to Dro compared to the wt strain (**Fig. 2a**). These results strongly suggest that the ribosome is the primary target of Dro in bacterial cells and that this PrAMP inhibits bacterial growth by interfering with translation termination.

**Fig. 2:**
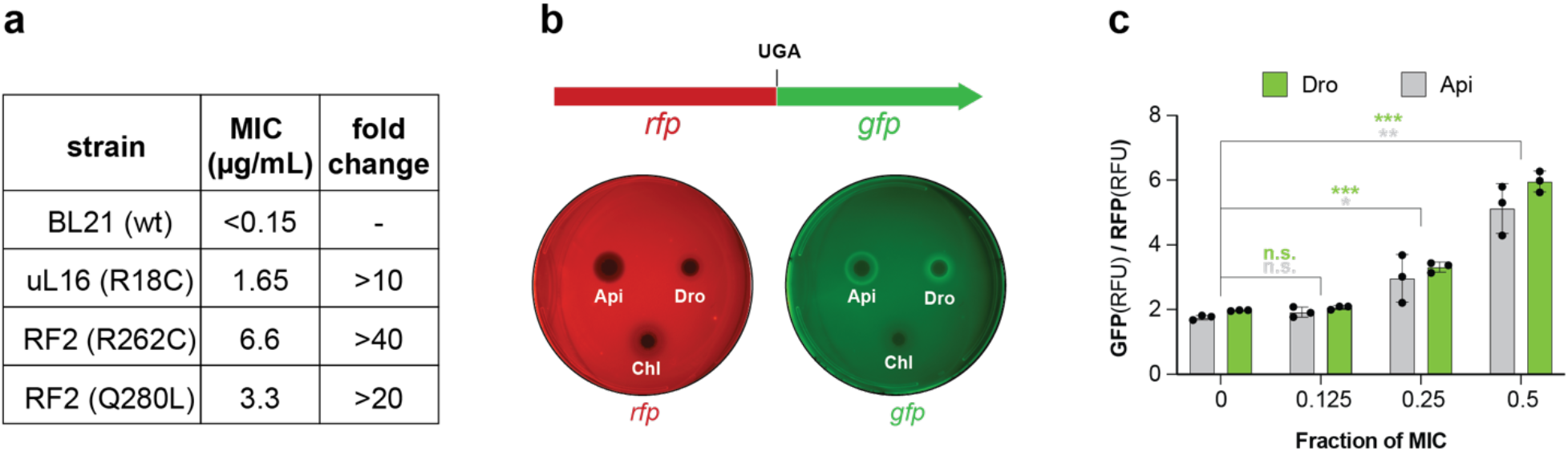
Antibacterial action of Dro involves the ribosome and RFs and induces stop codon readthrough. **a**, Minimal inhibitory concentrations (MICs) of Dro against wild-type (wt) or mutant *E. coli* BL21 cells transformed with the pSbmA plasmid. The mutant strains contained amino acid changes in ribosomal protein uL16 (Arg18Cys) or in type I RF2 (Arg262Cys or Gln280Leu). **b**, Top: Schematics of the *rfp-gfp* reporter in the pRXG plasmid ^44^. Expression of GFP depends on the readthrough of the in frame stop codon separating the two genes. Bottom: Dro-mediated readthrough activity determined by drop-diffusion test in *E. coli* BL21 cells carrying the pRXG[UGA] reporter plasmid. The green fluorescence halos on the cells plated on M9 minimal medium with agar supplemented with the reporter inducer IPTG indicate stop codon readthrough by Dro. The known readthrough activity mediated by Api137 (Api) ^25^ was used for comparison. Unrelated translation inhibitor chloramphenicol (Chl) was added as a negative control. The red plate demonstrates even expression of RFP on the reporter cells. Plate images were captured in the Cy3 (for RFP) and Cy2 (for GFP) channels of the imager and false-colored. **c**, Stop codon readthrough in liquid cultures. BL21 cells transformed with the pRXG[UGA] reporter plasmid were grown in liquid cultures in the presence of varying sub-inhibitory concentrations of Dro or Api with continuous monitoring of light absorbance (A_600_) and fluorescence of RFP and GFP proteins. The bar graph represents the ratios of GFP and RFP fluorescence values after 9 h of culture growth. For this experiment, the concentrations of Dro or Api at which no growth was observed at 9 h were taken as MIC and was 1 μM for Api and 4 μM for Dro. Statistical significance of difference from the no PrAMP control is shown (2-tailed *t*-test: n.s., not significant, p > 0.05; * 0.01< p ≤ 0.05; ** 0.001 < p ≤ 0.01; *** p ≤ 0.001).

### Dro induces stop codon readthrough by sequestering RFs

Decreased efficiency or availability of class-1 RFs stimulates stop codon readthrough ^43^. For instance, Api-mediated sequestration of RFs on a small fraction of cellular ribosomes prevents the rest of the ribosomes from releasing the nascent proteins at stop codons, leading to the eventual stop codon bypass by amino acid misincorporation ^25,32^. To test whether Dro action in bacteria leads to stop codon readthrough, we employed the reporter in which in-frame fused *rfp* and *gfp* genes are separated by a UGA stop codon ^44^. After introducing the reporter plasmid in *E. coli* BL21 cells and plating lawn of cells onto agar plates, we tested whether addition of PrAMP would stimulate GFP expression. At subinhibitory concentrations, both Dro and Api, but not the control antibiotic chloramphenicol (Chl), readily induced GFP expression (**Fig. 2b**). Similar results were obtained in the liquid culture experiments where cells carrying the readthrough reporter plasmid were exposed to varying sub-MIC concentrations of Dro and Api (**Fig. 2c**). Because neither Dro nor Api increased expression of the reference RFP reporter (**Fig. 2b**), this result means that both PrAMPs greatly stimulate stop codon readthrough thereby revealing that similar to Api, Dro likely sequesters the RFs on the terminating ribosome.

### Endogenously expressed Dro retains its mechanism of action and prevents bacterial growth

Having established the mechanism of Dro action, we proceeded to define the structural features of this PrAMP critical for its ability to inhibit translation termination. Previous efforts to improve the antibiotic activity of Dro involved introduction of specific amino acid changes in its structure ^33,45,46^. Some of these alterations indeed yielded derivatives with somewhat improved antibacterial properties or serum stability ^45,47-49^. However, labor- and time-consuming peptide synthesis limits the systematic evaluation of the contribution of the PrAMPs residues for activity. Furthermore, testing synthetic PrAMPs in whole-cell assays makes it challenging to evaluate whether the structural alterations affect cellular uptake and/or on-target activity.

Because our biochemical and microbiological experiments have verified high activity of the non-glycosylated Dro, we decided to evaluate the structure-activity relationship (SAR) of Dro by expressing the mutant variants directly in the bacterial cell via translation of plasmid-borne *dro* genes. This approach, which has been successfully used for the analysis of several other PrAMPs ^11,50-55^, allows bypassing the need for import into the cell and therefore directly reveals the contribution of individual amino acid residues to the action of the PrAMP upon the ribosome.

SAR studies of the in vivo-expressed PrAMPs require that the endogenously produced peptide retains its antibacterial activity and inhibits cell growth by the same mechanism as the wt PrAMP prototype. To test for the inhibitory activity of endogenously expressed Dro, its coding sequence (equipped with start codon, UGA or UAG stop codons, and codon-optimized for expression in *E. coli*) was introduced into the pDro[UGA] or pDro[UAG] plasmids under control of the P_BAD_ promoter inducible by L-arabinose (L-Ara) ^11^ (**Fig. 3a**). Surprisingly, in contrast to the highly toxic expression of Api ^11^, L-Ara-mediated expression of Dro did not prevent the growth of the plasmid-transformed *E. coli* BL21 cells on LB/agar plates (**Extended Data Fig. 2**). Nevertheless, because our in vitro assays hinted that the activity of Dro seemed to be somewhat weaker than that of Api (**Fig. 1c,d**), we reasoned that toxicity of Dro could be more pronounced in cells growing under less favorable conditions. Indeed, growth in a minimal (M9) medium rendered *E. coli* BL21 cells susceptible to the endogenous expression of Dro (**Fig. 4**).

**Fig. 3:**
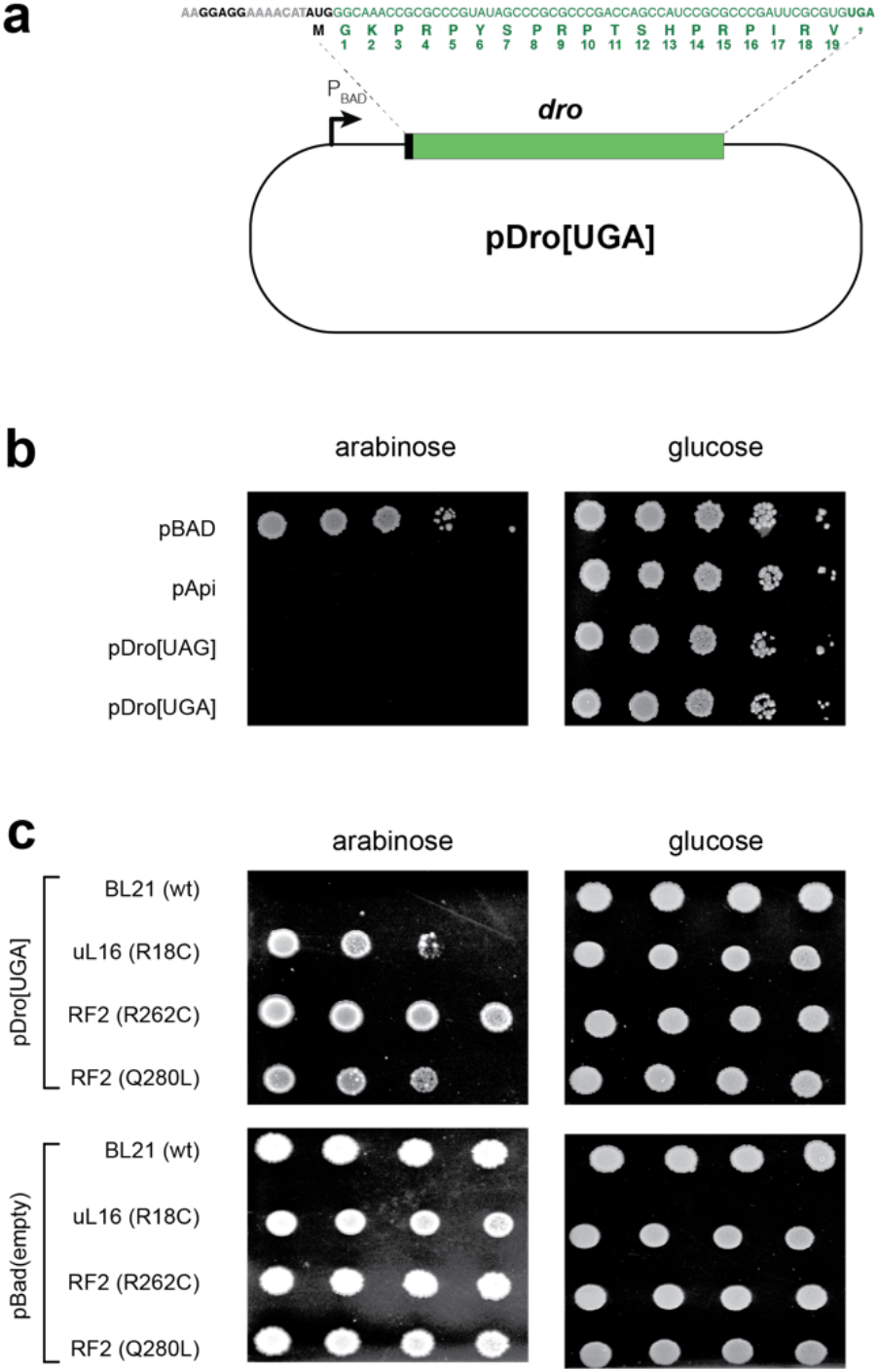
Endogenous expression of Dro is toxic for bacteria due to the interactions of the PrAMP with the ribosome and release factors. **a**, Schematics of the pDro[UGA] plasmid for expression of Dro in *E. coli* cells. The L-arabinose inducible P_BAD_ promoter is indicated. The Dro encoding sequence, preceded by a ribosome binding site and a start codon, is shown in green. The start and stop codons are in bold. Numbering of the Dro residues, corresponding to the naturally produced PrAMP, is shown below the amino acid sequence. The sequence of the pDro[UAG] plasmid (not shown) is identical except that the stop codon of the *dro* ORF is UAG. **b**,**c**, Spot test to determine the effect of endogenous Dro expression in (**b**) wt *E. coli* BL21 cells or (**c**) BL21 derivatives carrying mutations in ribosomal protein uL16 or RF2. Ten-fold (panel **b**) or three-fold (panel **c**) dilutions of cell cultures were spotted on M9 minimal medium agar plates under non-inducing (glucose) or inducing (arabinose) conditions. Api-expressing cells (pApi) were used for comparison. Cells transformed with an empty pBad vector were used as control. The effect of Dro expression upon cells grown in rich medium is shown in **Extended Data Fig. 3**.

**Fig. 4:**
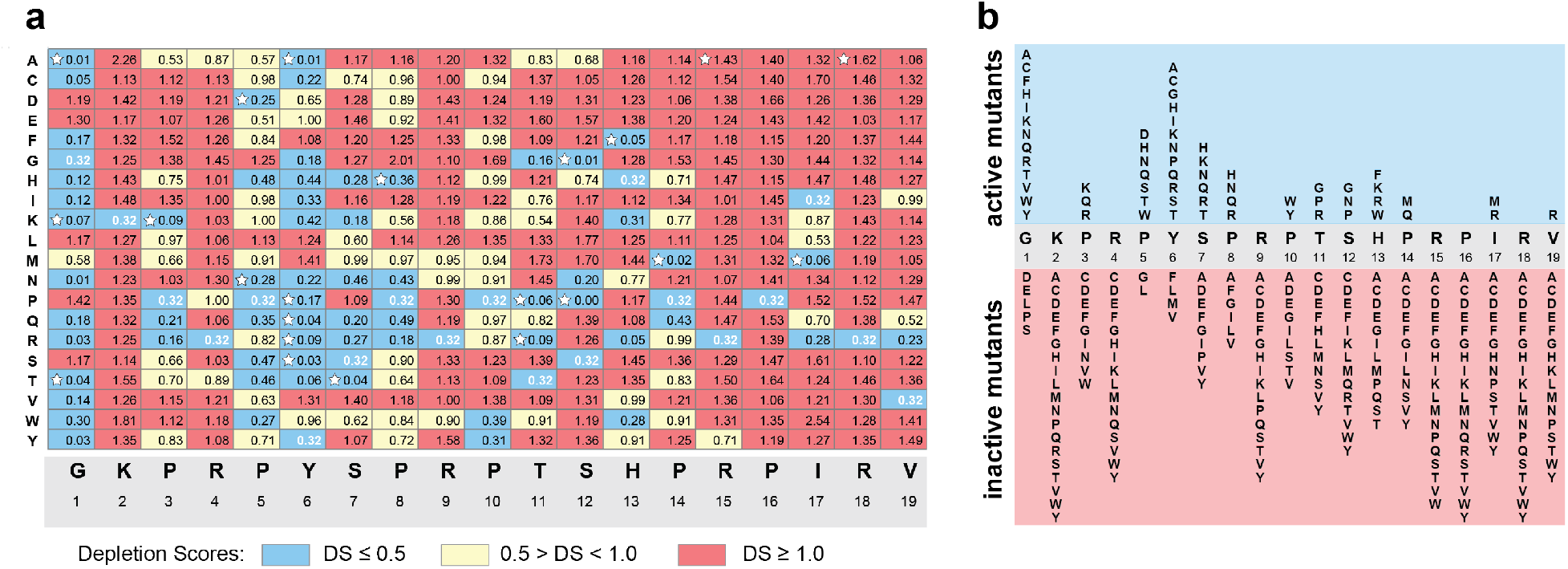
The growth inhibition effect of Dro mutants expressed in cells. **a**, Depletion scores (DS) characterizing effects of single-amino acid substitutions upon the ability of endogenously expressed Dro variants to arrest cell growth. DS reflects the difference in abundance of a particular variant in the library of mutant *dro* genes in pDro[UAG] plasmids expressed in *E. coli* BL21 cells grown under inducing versus non-inducing conditions. Dro mutants were classified as highly toxic (DS ≤0.5) (blue), mildly toxic (DS 0.5 to 1.0) (yellow) or not toxic (DS ≥1) (red). DS value (0.32) of the wt Dro is shown with white numbers. Variants selected for chemical synthesis and further analysis are marked with white stars. The DS plot of the mutant library in pDro[UGA] plasmids is shown in **Extended Data Fig. 4b. b**, Diagram summarizing the effect of the Dro mutations on cell growth. Only highly toxic (blue field) and non-toxic (red field) mutants are shown.

Because the expression of a foreign protein could be inhibitory to the cell for a variety of reasons ^56^, it was important to verify that the toxicity of endogenously expressed Dro is due to its ability to interfere with translation termination. To this end, we once again exploited the Api- and, as we now know, Dro-resistant mutants with alterations in the ribosome (protein uL16) or in RF2 ^25^ (**Fig. 2a**). We found that Dro-resistant mutants transformed with the pDro plasmid tolerated the endogenous expression of Dro much better than the parental wt strain (**Fig. 3c**). These results demonstrate that when translated in the bacterial cell, Dro retains its ability to bind to the ribosome and interfere with the termination of translation.

### Screening of comprehensive Dro mutant libraries reveals residues critical for the on-target activity

Although Dro and Api show a convergent mode of action, they have significant differences in their amino acid sequences (**Fig. 1b**). Therefore, it was important to deduce the contribution of individual residues of Dro to ribosome binding, inhibition of translation, and cell growth. To achieve this goal, we assessed the on-target activity of each single amino acid mutant of Dro. Having verified that the endogenously expressed Dro retains its mode of action, we prepared two libraries of mutant Dro genes, with ORFs ending with either UAG or UGA stop codons, and analyzed the inhibitory activity of the endogenously expressed Dro mutants. Each library encoded all possible single amino acid Dro mutants (n=361) plus the wt. After introducing the mutant genes in the original pDro plasmid, the libraries were transformed into *E. coli* BL21 cells and plated at high density onto M9/agar plates supplemented with glucose (non-inducing conditions) or with L-Ara (inducing conditions). After deep sequencing of the libraries, we computed the ‘depletion score’ (DS) for each mutant that reflected the difference in its abundance in the libraries grown under inducing vs non-inducing conditions. Clones expressing inhibitory peptides are expected to be depleted in the inducing (L-Ara) conditions and the DS values of such mutants should be less than 1. Conversely, the relative abundance of clones expressing inactive Dro variants should increase on the L-Ara plate and the respective DS values should be above 1 (**Fig. 4**). Consistent with this metric, the DS value of wt Dro is 0.3. The results of the analysis of the two libraries (ending with UGA or UAG) were highly convergent (Pearson correlation coefficient of 0.91) (**Fig. 3a,b**, **Extended Data Fig. 4a**,**b**), indicating the independence of the activity of the in situ translated Dro peptides from the nature of the stop codon of the *dro* gene and emphasizing the robustness of the obtained results.

The overall distribution of Dro-inactivating mutations shows a characteristic bias with many of them clustering towards the C-terminal segment of the PrAMP (**Fig. 4; Extended Data Fig. 3b**, chart cells colored red). In particular, the sequence encompassing Thr11-Val19 appears to be critical for the on-target activity, as most mutations of these residues inactivate the endogenously expressed Dro. Three additional residues in the N-terminal segment of Dro, Lys2, Arg4, and Arg9, are also pivotal for the activity: any mutation of these positively charged amino acids reduce the activity of the PrAMP. In contrast, the Pro5-Pro8 segment and the N-terminal glycine can tolerate multiple mutations without any significant activity loss (**Fig. 4; Extended Data Fig. 3b**, chart cells colored blue). Thus, our mutational data strongly argue that an extended C-terminal segment of Dro as well as some of its residues proximal to the N-terminus are important for binding of the PrAMP to the ribosome and trapping the RF (**Extended Data Fig. 3c**). This pattern contrasts the results of similar mutational studies of the other type II PrAMP, Api, where only a few C-terminal amino acid residues were found to be functionally critical, whereas the N-terminal sequence could be mutated or even deleted without significant impact on its activity ^11^. Thus, while sharing the general mechanism of action, different type II PrAMPs utilize converging but clearly distinct strategies for target engagement.

### Improved activity of specific Dro variants

Remarkably, some of the endogenously expressed Dro mutants yielded DS values lower than that of the wt PrAMP, suggesting that they may be more active than the wt non-glycosylated Dro. To directly explore this possibility, we chemically synthesized the potentially improved Dro variants to test their activity both in vitro and in vivo. Twenty synthetic peptides carrying mutations at different Dro residues expected to preserve or possibly improve the activity were prepared (**Fig. 4a**, white stars). In addition, we also generated two mutants, R_15_A and R_18_A, that could serve as negative controls because their DS values of 1.43 and 1.62, respectively, indicate that they are likely inactive variants (**Fig. 4a**).

Toeprinting assays showed that the synthetic Dro variants selected as potentially active are able to stall the ribosome at the UAG stop codon of the model ORF, while those identified as likely inactive (the R_15_A and R_18_A mutants) failed to do so (**Extended Data Fig. 4**). In agreement with the activity charts (**Fig. 4; Extended Data Fig. 3b**), several of the synthetic peptides, including the S7T, T11P, T11R, S12G, H13F, P14M and I17M mutants, arrested translation more efficiently than the unmodified Dro (**Fig. 5a**). Strikingly, three of the tested mutants, T11P, T11R and particularly P14M, stalled the ribosome even better than Api137, which so far is known to be the most active type II PrAMP ^11,25,30^. These results suggest that the ribosome-inhibitory activity of non-glycosylated Dro can be significantly improved by small alterations in the peptide sequence.

**Fig. 5:**
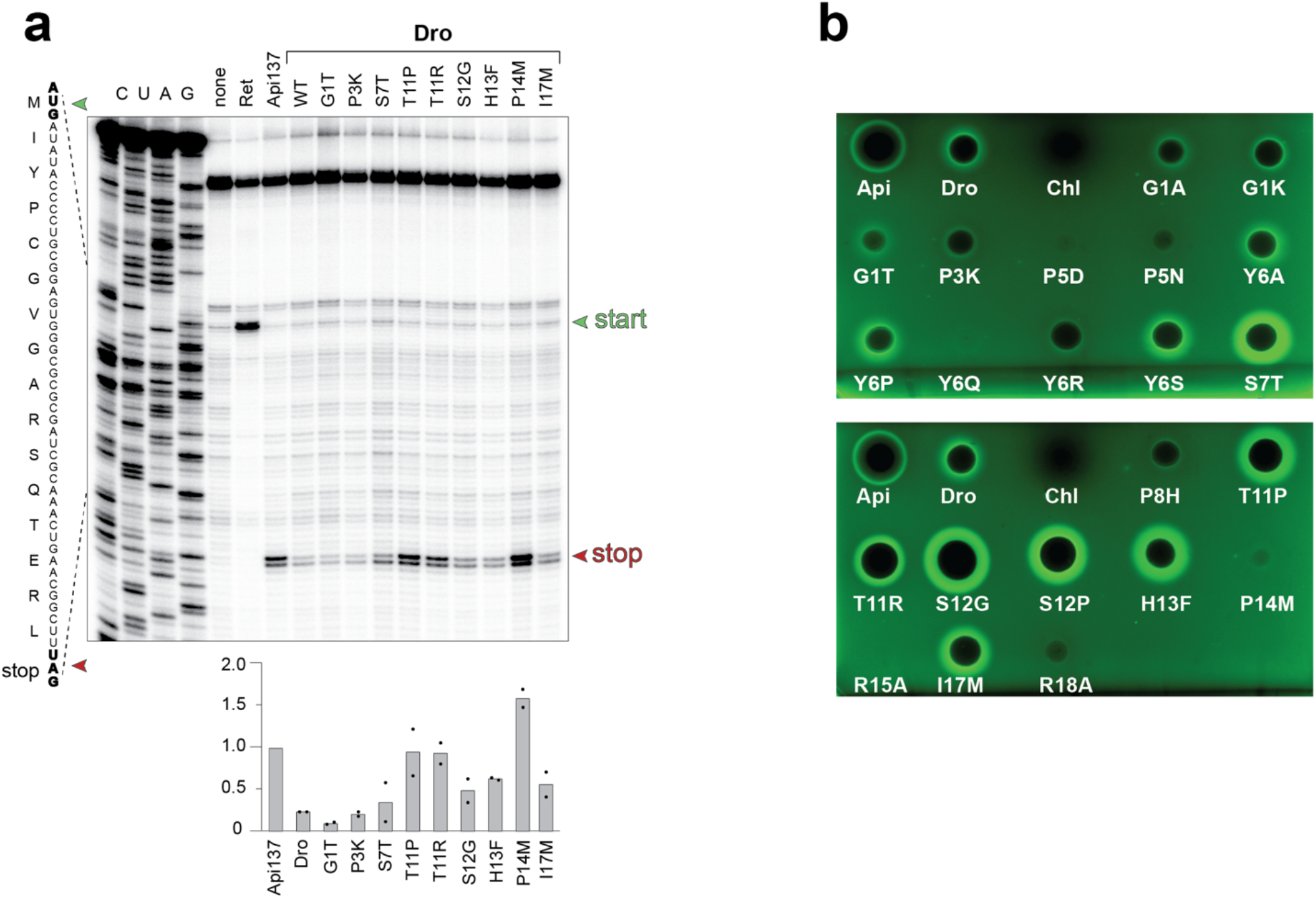
Synthetic Dro variants with single-amino acid substitutions preserve their ability to interfere with translation termination. **a**, Toeprinting analysis of the ribosome arrest at the stop codon (red arrowhead) of the model *yrbA* ORF caused by synthetic wt or mutant non-glycosylated Dro variants. Arrest caused by Api137 was used as a positive control. The antibiotic retapamulin (Ret) was used to mark the toeprint band generated by ribosomes arrested at the start codon (green arrowhead). Bar graph shows the normalized average intensity of the toeprint bands of ribosomes arrested at the stop codon estimated from two independent experiments. Individual values from the two experiments are marked with dots. Intensity of the toeprint band generated by Api137-arrested ribosomes was set as 1.0. The complete toeprinting analysis of all the synthesized mutants is shown in **Extended Data Fig. 4. b**, Antibacterial (the clearing zones) and stop codon readthrough (the GFP fluorescence halos) activities of synthetic Dro variants in *E. coli* BL21 cells transformed with the pRXG[UGA] reporter plasmid. Api137 (1/10 amount relative to Dro variants) was used for comparison and Chl was used as a stop codon read-through negative control.

The antibacterial activity of the PrAMPs requires their efficient cellular uptake, which is facilitated by the SbmA-type peptide transporters ^26,27^ Because in our library screening experiments the Dro variants were expressed inside the cell, it was unclear whether any of the PrAMPs, including those with the improved on-target activity, would exhibit antibacterial properties when added exogenously to cells. Therefore, we tested the antibacterial action of our synthetic Dro variants by determining their MICs against *E. coli* BL21 cells (**Extended Data Table 1**). The MIC values of most of the mutants were comparable to those of the non-glycosylated wt Dro demonstrating their antibiotic activity. Interestingly, the MIC of one of the negative control mutants, R18A, was only 4 times higher than that of the wt Dro control (2 μg/mL), indicating that this PrAMP variant retained some antibacterial activity against *E. coli*. In comparison, the other negative control mutant, R15A, showed a very weak activity with an MIC value of >64 μg/mL.

We further tested whether the synthetic Dro variants were able to induce stop codon readthrough. We used the drop diffusion assay with the *E. coli* BL21 cells carrying the RFP-GFP reporter (**Fig. 2b**) to simultaneously verify the cell-inhibitory activity of Dro mutants and test their ability to induce stop codon bypass. In agreement with the MIC results, the majority of the tested mutants inhibited *E. coli* growth on agar plates and most of the inhibitory mutants were able to induce stop codon readthrough (**Fig. 5b, Extended Data Table 1**). The extent of GFP induction, which requires translation through the stop codon (**Fig. 2b**), generally matched with the ability of the Dro variants to stall the ribosome at the stop codon in the in vitro toeprinting assay (**Fig. 5a**). Thus, consistent with the findings of the in vitro experiments, the mutants S7T, T11R, T11P, S12G induced stop codon readthrough in vivo much more efficiently than wt Dro (**Fig 5a, b**).

Altogether, our data show that by introducing limited mutations into the structure of the unmodified Dro it is possible to significantly improve target engagement and possibly antimicrobial activity of this PrAMP without requiring synthetically challenging posttranslational glycosylation.

## Discussion

In this work we examined the mechanism of action of the PrAMP Dro encoded in the fruit fly genome. We found that Dro interferes with bacterial growth by acting upon the ribosome and arresting it at stop codons likely trapping class 1 RFs in the post-release complex. Interrogating comprehensive libraries of single amino acid mutants, we identified Dro residues critical for the PrAMP’s ability to inhibit translation. We further demonstrated that specific mutations could increase the ability of non-glycosylated Dro to arrest the terminating ribosome, stimulate stop codon readthrough and possibly improve the antibacterial properties.

The mechanism of Dro action resembles that of the only known type II PrAMP, Api. However, as our mutational analysis shows, Api and Dro employ notably different strategies for target engagement. Api requires only few of its C-terminal residues to bind tightly to the ribosome; mutations of most of its other amino acids do not abolish Api’s ability to interfere with translation and with cell growth ^11^ (**Extended Data Fig. 3c**). Consistently, in the cryo-EM reconstructions of the ribosome-Api137 complex, it was observed that the peptide’s N-terminal segment is disordered, indicating that it forms only weak non-specific interactions with the NPET ^25^. In contrast, mutations of most of Dro residues lead to activity loss, indicating that this PrAMP establishes a more extensive network of critical interactions with the ribosome. Functionally important amino acid residues are not confined to the C-terminal portion of Dro sequence, but rather distributed through the entire length of the PrAMP and some of them (Lys2, Arg4, and Arg9) are found in the Dro’s N-terminal segment. This finding is consistent with the cryo-EM structure of the ribosome-Dro complex (see the accompanying paper of Koller et al. ^57^), where the entire peptide is well resolved due to the multiple interactions with the NPET. Our mutational data help to discern which of the contacts between Dro residues and the ribosome observed in the cryo-EM reconstructions of Koller at al. ^57^ are essential for the PrAMP’s activity and which may play only a secondary role (**Extended Data Fig. 5**). Every mutation of the four Arg residues of Dro (Arg4, Arg9, Arg15, and Arg18) diminishes activity, indicating their importance for Dro action. Most likely stacking of Arg9 side chain upon the base of A751 and of Arg15 upon A2062 of the 23S rRNA and the hydrogen bonding of Arg9 with the ribosomal protein uL22 and of Arg18 with RF, are pivotal for Dro action (**Extended Data Fig. 5a-c**). The rationale for the importance of Arg4 is less clear because in the cryo-EM reconstructions of the ribosome complexed with the Thr11-glycosylated Dro, its closest interacting partner, the non-bridging oxygen of the U1258 phosphate, is 4 Å away – too far for a strong hydrogen bond (**Extended Data Fig. 5d**). It is possible, however, that the lack of Thr11 glycosylation may cause a minimal relocation of Arg4 in the NPET strengthening its interaction with U1258 making this contact particularly important for the non-glycosylated Dro. Two other Dro residues that do not tolerate any substitution are Lys2 and Pro16 (**Fig. 4**). Functional significance of Lys2 may indicate that its interaction with the G1256 phosphate is functionally important (**Extended Data Fig. 5e**). In contrast, Pro16 does not engage in direct interactions with the ribosome; its importance likely stems from the necessity to properly orient the neighboring Arg15 for its contacts with A2062 and/or to position the peptide’s C-terminus for interactions with the RF (**Extended Data Fig. 5a**,**b**).

Conversely to the aforementioned critical residues of Dro, the Pro5-Pro8 segment can tolerate multiple mutations (**Fig. 4**). While in the cryo-EM model Tyr6 and Ser7 form direct contacts with the ribosomal proteins uL4 and uL22 at the NPET constriction (**Extended Data Fig. 5f**), such interactions either contribute insignificantly to the Dro binding or can be replaced with alternative interactions provided by the mutated residues. This conclusion is consistent with the results of the previous limited mutational studies where several tested substitutions of Tyr6 or Ser7 had negligible effect upon activity of Dro ^47^.

The presence of N-acetylglucosamine sugar at Thr11 of the native Dro could potentially affect the exact placement of the PrAMP in the NPET. It is possible, therefore, that some of the identified mutations that enhance potency of the unmodified Dro compensate for the lack of the Thr11 modification. Thus, Pro14 closely approaches the N-acetylglucosamine of the ribosome-bound native Dro, potentially facilitating the interaction of the sugar residue with 23S rRNA U2609. The conformational constraint imposed by the rigid proline residue upon Dro’s trajectory, beneficial for binding of the glycosylated PrAMP, may become suboptimal when the sugar is absent. The replacement of Pro14 with methionine, which significantly stimulates the ribosome-arresting ability of non-glycosylated Dro (**Fig. 5a; Extended Data Fig. 4**) might relax the proline-imposed structural constraints allowing for additional stabilizing contacts, possibly involving the methionine’s side chain.

While illuminating, the results of our in vivo mutational analysis should be interpreted with a certain degree of caution. The activity of the endogenously expressed Dro could depend on its stability, which can be affected by the mutations. It is also possible that alterations of the early codons of the *dro* gene may affect the efficiency of Dro expression rather than activity. Thus, most of the Gly1 codon substitutions seem to yield peptides more active than wt Dro (**Fig. 4**). However, in the toeprinting experiments, readthrough assays or MIC testing, none of the three tested synthetic peptides with Gly1 mutations performed any better than wt Dro (**Fig. 5; Extended Data Fig. 4**). By the same token, the inactivating mutations of the Lys2 codon could potentially diminish expression of the *dro* gene. Regardless of these cautionary notes, it remains highly likely that survivability of most of the *E. coli* library clones endogenously expressing Dro variants directly reflects the ability of the PrAMP variants to bind to the ribosome and inhibit translation.

Despite the sequence divergence and significantly different modes of interaction of Dro and Api with the ribosome, binding of these distinct peptides leads to formation of comparable complexes. These complexes strikingly resemble the structure of the terminating ribosome immediately after the RF-promoted hydrolysis of peptidyl-tRNA, where the type II PrAMPs mimic the orientation and placement of a newly hydrolyzed nascent protein. While completed nascent proteins rapidly dissociate from the ribosome, Dro and Api remain stably bound due to their interactions with the NPET. Conceivably, endogenously expressed type II PrAMPs may remain stuck in the NPET of the ribosome on which they were just synthesized. The possibility that wt Api, non-glycosylated Dro, and many of their multiple variants can remain within the NPET after their synthesis, suggests that the C-terminal segment of a wide spectrum of proteins may form strong enough interactions with the NPET to allow for the nascent protein to linger in the ribosome after separation from tRNA. Identifying such sequences in the cellular proteome could provide new insights into translation regulation, protein folding and targeting. The structural malleability of the type II PrAMPs also opens broad opportunities for optimizing their sequence and structure for medical applications.

Our in vitro and in vivo data strongly argue that inhibition of translation termination likely involving trapping of RFs on the ribosome is the primary mode of action of type II PrAMPs, including Dro. However, this view may not capture the entire spectrum of the effects produced by these inhibitors in the bacterial cell, which may differ between individual Dro mutants. For example, we have noted that the Thr11Arg mutant was more active than the His13Phe variant in arresting the ribosome at the stop codon in vitro (**Fig. 5a**), whereas the latter generated a broader and brighter GFP fluorescence halo in the in vivo drop-diffusion readthrough assay (**Fig 5b**). Similarly, the synthetic Tyr6Ser and Ser7Thr or Thr11Arg and Ser12Gly Dro mutants showed comparable antibacterial activity, yet at the subinhibitory level of endogenous expression, the Ser7Thr variant was more potent than Tyr6Ser and Ser12Gly mutant was more potent than Thr11Arg in stimulating GFP expression via stop codon readthrough (**Fig. 5b**). Possibly additional, more cryptic modes of action, e.g., inhibition of initiation of translation of some genes, as was observed for Api ^32^, could contribute to the antibacterial action of some of the Dro mutants.

The antibacterial action of the PrAMPs relies on their ability to reach the cytoplasmic target. Dro is imported into *E. coli* cells by SbmA transporter whose specificity is unknown ^26,27^. Some mutations that stimulate the on-target action of endogenously expressed Dro may at the same time diminish its intracellular accumulation when the PrAMP is added exogenously. Indeed, while the mutation of Pro14Met renders Dro more active in the cell-free system (**Fig. 5a**), it dramatically reduced its antibacterial properties (**Fig. 5b; Extended Data Table 1**) because the mutation likely prevents uptake. Conversely, while the Arg18Ala substitution abolished the ability to stall the ribosome (**Extended Data Fig. 4**) or induced stop codon readthrough (**Fig. 5b**), the mutations did not fully prevent the PrAMP from inhibiting bacterial growth (**Fig. 5b; Extended Data Table 1**). It is conceivable that the Arg18Ala mutation alters the mode of the PrAMP’s action upon the ribosome or maybe even allows engagement of a different cellular target.

In conclusion, our study has not only revealed the mode of action of Dro and identified its amino acid residues critical for activity, but it also showed that by substituting only a few amino acids in the structure of a non-glycosylated PrAMP, it is possible to improve its on-target activity and possibly antibiotic properties. This may allow further optimization of Dro and related PrAMPs for potential therapeutic use bypassing the necessity for the experimentally challenging introduction of posttranslational modifications.

## Data availability

The uncropped gels and the raw data used for preparing charts and graphs are shown in the Source Data files associated with the manuscript.

## Author contributions

RH, NV-L and ASM conceived the study; KM guided and supervised preparation of the endogenously-expressed wt Dro and Dro mutant library; DK carried out toeprinting and microbiological experiments; AB and AK synthesized peptides and carried out in vitro translation and MIC testing experiments; IO cloned Dro gene and carried out library screening experiments; KM and CB analyzed library screening results; CB consulted on multiple experiments; WH analyzed stop codon readthrough and Dro-resistant mutants; YP analyzed structural data; KM, CB, AK, RH, NV-L and ASM analyzed data; KM, NV-L and ASM wrote the manuscript.

## Acknowledgements

We thank Timm Koller and Daniel Wilson (University of Hamburg) for generously sharing their results and the structure of the ribosome-drosocin complex. This work was supported in part by NIH grant NIAID R01AI162961 (to ASM, NV-L and YSP).

## Methods

### Peptides and oligonucleotides

Apidaecin 1b (Api1b) was synthesized by Genescript and Api137 was synthesized by NovoPro Biosciences, Inc., both at 95% purity. All the other peptides were synthesized in-house on solid phase as described ^38^. The DNA oligonucleotides libraries for generating Dro mutant libraries were ordered from Twist Biosciences. All other DNA oligonucleotides were from Integrated DNA Technologies.

### In vitro translation and toeprinting analysis

The sf-GFP reporter protein was expressed essentially as described ^38^ in the PURExpress ΔRF123 system (New England Biolabs). The DNA template was PCR-amplified from the pY71sfGFP plasmid ^58^ by PCR using primers sfGFP-fwd and sfGFP-rev (all primers are listed in Supplementary Table S2). Transcription-translation reactions containing 35 ng of the DNA template were carried in a final volume of 5 μL. When needed, 0.5 μL of a 1:50 dilution of the RF1solution included in the kit were added to the reaction. Fluorescence of sfGFP was recorded in a microplate reader (Gemini EM, Molecular Devices) at 37°C for 2 h (λexc = 485 nm, λem = 535 nm). Fluorescence values used for **Extended Data Fig. 1a** correspond to the time point of 2 h.

Toeprinting experiments were carried out following the procedure described previously ^25,59^. The *yrbA-fs15* DNA templates (with TAG or TGA stop codons) used for in vitro transcription/translation reactions in the classic PURExpress system (New England Biolabs) were prepared by PCR, as previously described ^11,25^, by combining the overlapping primers T7, T7-IR-AUG, IR-yrbA-fs15-RF1 (for TAG) or IR-yrbA-fs15-RF2 (for TAG), post-NV1, and NV1, rendering the sequences TAATACGACTCACTATAGGGCTTAAGTATAAGGAGGAAAACAT<u>ATGATATACCCCT GCGGAGTGGGCGCGCGATCGCAAACTGAACGGCTT(</u>**<u>TAG</u>**<u>/</u>**<u>TGA</u>**)GCCGACCTCGAC AGTTGGATTCACGTGCTGAATCCTGATGCGATGTCGAGTTAATAAGCAAAATTCATTATAACC (ORF sequence is underlined). PrAMPs or retapamulin were added from stock solutions in water to a final concentration of 50 μM.

### MIC testing

Antimicrobial activity of Dro or its variants was determined by the MIC test in 96-well plates as described ^11,38^. Tests were performed with wild-type *E. coli* BL21(DE3) cells transformed with the pSbmA plasmid or its derivatives containing mutations in uL16 (Arg81Cys) or in RF2 (Arg262Cys or Gln280Leu) ^11^. MIC testing of synthetic Dro variants was carried out using untransformed BL21(DE3) cells. Testing the effects of *dnaK* knock-out was carried out in BW25113 strain or its *dnaK::kan* derivative from the Keio collection ^60^. Tests were performed in cell cultures grown in 25% Cation-adjusted MHB medium. PrAMPs were serially diluted from stock solutions in water.

### Stop codon readthrough assay

The ability of Dro or its variants to cause readthrough of premature stop codons in bacterial cells was tested by the dual RFP/GFP fluorescence pRXG plasmid reporter assay. *E. coli* BL21 cells were transformed with the original pRXG plasmid, where RFP and GFP encoding genes are separated by the TAG stop codon ^44^, or an engineered mutant version of the original pRXG plasmid where the stop codon was changed from TAG to TGA. Cell were grown overnight in LB medium supplemented with 50 μg/mL kanamycin (Kan) and then diluted 1:10 into fresh LB/Kan medium. Exponentially growing cells at an A_600_ of ∼1 were pelleted and resuspended in the same volume of M9 minimal medium supplemented with 2 mM MgSO_4_, 0.1 mM CaCl_2_, 10 μg/mL thiamine. The cell suspension (3 mL) was poured over an agar plated prepared on supplemented M9 minimal medium and additionally containing 0.2 mM IPTG and 50 μg/mL Kan. After a brief (∼30 s) incubation, excess liquid was aspirated, and plates were allowed to dry with lids removed. Solutions of Dro or its derivatives (2 mM), of Api137 (0.2 mM) (2 μL drops) or of 1 mg/mL of Chl (1 μL drop) were applied. Plates were incubated for 18 to 24 h at 37°C and imaged in a ChemiDoc Imaging System (BioRad) to detect RFP and GFP fluorescence (Cy3 and Cy2 channels, respectively). Images were processed and false-colored using the ImageLab software (Bio-Rad).

For the analysis of stop codon readthrough in liquid culture, *E. coli* BL21 cells transformed with pRXG[UGA] plasmid were gown at 37°C overnight in LB medium supplemented with 50 μg/mL Kan, then diluted into the same medium and grown to A_600_ ∼0.4. Cultures were then diluted 1:100 into M9 medium supplemented with 0.4% glucose and 0.2 mM of IPTG, placed into wells of a 96-well (black-well clear-bottom) plate and varying concentrations of Dro or Api were added. The final culture volume was 100 μL per well. The PrAMP concentrations varied from 0 to 16 μM over two-fold dilutions. Cells were grown in the TECAN plate reader at 37 °C with monitoring every 20 min of cell density (A_600_), RFP fluorescence (λ ex = 550 nm, λ em = 675 nm, gain =100) and GFP fluorescence (λ ex = 485 nm, λ em = 520 nm, gain =125). Data were plotted and data points corresponding to 9 h of growth were used for calculating the GFP/RFP (RFU/RFU) ratios.

### Endogenous expression of Dro in bacterial cells

The sequence of the DNA templates (synthesized by IDT) coding for the mature Dro peptide (GKPRPYSPRPTSHPRPIRV) was codon-optimized for expression in *E. coli* and ATG start codon and stop codons TAG or TGA (underlined in the sequence provided below) were added: ATGGGCAAACCGCGCCCGTATAGCCCGCGCCCGACCAGCCATCCGCGCCCGATT CGCGTG<u>TAG</u>(or <u>TGA</u>) The Dro-encoding templates were used to swap the Api encoding gene with the Dro sequence in the pApi vector ^11^. For this, the Dro templates were introduced by Gibson assembly into pApi cut with SacI and XbaI restriction enzymes, yielding pDro[UAG] and pDro[UGA] plasmids.

To test the effects of the endogenously expressed Dro on cell growth, the pDro[UAG] and pDro[UGA] plasmids were transformed into *E. coli* BL21 cells. Transformants were grown overnight in LB medium supplemented with 30 μg/ml Chl and 2% glucose. Cultures were diluted 100-fold into fresh medium, grown at 37°C, and upon reaching an A_600_ ∼0.5 they were spun and resuspended in LB or M9 medium. Ten-fold serial dilutions were spotted (3 μl) on LB/agar or supplemented M9/agar plates containing 30 μg/ml Chl, and either 2% glucose (non-inducing conditions) or 2% L-Ara (inducing conditions). The plates were incubated at 37°C for 16-48 h and photographed.

### Spot-test

*E. coli* cells (BL21 or the mutant strains) transformed with pDro or pApi plasmids or with the empty pBad vector were grown overnight in LB medium supplemented with 30 μg/mL Chl and 0.4% glucose. Cultures were diluted 1:100 into M9 medium supplemented with 2% glycerol, 2 g/L tryptone and 30 μg/mL Chl and grown to 10^7^ – 10^8^ cfu/mL. Three μL of ten-fold (Fig. 3b) or three-fold (Fig. 3c) dilutions were spotted on M9 plates supplemented with 2% glycerol, 2 g/L tryptone, 30 μg/mL Chl, and containing either 0.4% glucose or 0.2 % L-arabinose. Plates were grown 36-48 h and photographed.

### Mutant Dro library construction and selection screening

#### Generating plasmid libraries

Two libraries of the Dro genes, one ending with the TGA stop codon and another ending with the TAG codon were generated by massively parallel oligonucleotide synthesis. Each of the codons of the Dro-encoding gene was individually replaced by one of the 19 codons (with the highest *E. coli* codon adaptation index value) specifying each of the alternative amino acid residues. The dsDNA fragments (wt + 361 mutant for each library) of 363 nts each were procured from Twist Bioscience. The libraries were cloned into SacI/XbaI cut pBAD vector via Gibson Assembly, transformed into XL1-blue cells and plated on LB/agar plates supplemented with 30 μg/ml of Chl and 2% glucose yielding ∼4×10^5^ clones per library (>100-fold coverage). The clones were washed off the plates and the total plasmid was extracted using High-pure plasmid isolation kit (Roche).

#### Library selection

The amplified plasmid libraries were then transformed into *E. coli* BL21 cells by electroporation and plated on ⊘ 136 mm M9/agar plates containing 30 μg/mL Chl, 1% glycerol, and either 2% glucose or 2% L-Ara. The concentration of L-Ara used in the selection experiment corresponded to that required for strong inhibition of the growth of cells transformed with the original pDro[UAG] plasmid. Plates were incubated at 37° for 48 h. The clones (∼4×10^5^ per condition) were washed off the plates with 15% (v/v) glycerol, flash-frozen in liquid nitrogen and stored at −80°C.

#### Generating sequencing libraries

Frozen glycerol stocks of libraries washed off the glucose- or L-Ara plates were thawed, and total plasmid was isolated using High-pure plasmid isolation kit (Roche). The *dro* genes were minimally PCR amplified (10 cycles) using Dro-Lib-Fwd and Dro-Lib-Rev primers. The PCR products were isolated using a DNA Clean and Concentrator kit (Zymo Research). Another round of PCR amplification (10 cycles) with the primers IDT8_i57 – IDT8_512 and IDT8_i77 – IDT8_i712 was used to add Illumina sequencing adapters to the libraries from each condition. The PCR products were size selected in a 15% TBE-Urea gel followed by elution, ethanol precipitation, and quantification in a Qubit fluorometer. The libraries were spiked with 30% PhiX due to low complexity and sequenced on an Illumina MiSeq platform at the DNA sequencing facility of Northwestern University.

#### Sequence analysis and phenotypic scoring

Cutadapt (V3.4) was used to remove all the extra nucleotides from the 3’ and 5’ ends of the reads, leaving only the Dro sequence variants including the start and stop codons. Next, all the reads were sorted and counted based on unique sequence identity using the Unix command:

$ gzip -dc input_file.fastq.gz | paste - - - - | cut -f2 | sort | uniq -c | sort -g > output_file.txt Only exact matches to expected library sequences were considered for further analysis leaving a total of 432,013/179,563 sequences (glucose and arabinose conditions, respectively) for the UAG library and 163,449/295,374 sequences (glucose and L-Ara conditions, respectively) for the UGA library. The relative abundance of each mutant was calculated by the dividing the number of occurrences of its sequence by the total number of sequences in the library. Depletion score (DS) was then computed as log_2_ of the relative abundance of the variant sequence in the induced (L-Ara) sample divided by relative abundance of the same variant in the uninduced (glucose) samples. Further binning was as follows: DS values ≤ 0.5 - active Dro mutants; DS values > 0.5 but < 1.0 – moderately active mutants; DS values ≥ 1.0 – inactive mutants. The DS charts (**Fig. 4; Extended Data Fig. 3**) were generated in Excel.

