## Extended Data for "Inhibition of translation termination by Drosocin, an antimicrobial peptide from fruit flies"

| Dro mutants | MIC [µg/mL] | clearing zone (mm) |
|---|---|---|
| WT | 2 | 5 |
| G1A | 2 | 4 |
| G1K | 2 | 6 |
| G1T | 8 | <2 |
| P3K | 4 | 4 |
| P5D | 8 | - |
| P5N | 8 | <2 |
| Y6A | 4-8 | 5 |
| Y6P | 4 | 4 |
| Y6Q | 8 | - |
| Y6R | 2 | 6 |
| Y6S | 2-4 | 5 |
| S7T | 2 | 7 |
| P8H | 2 | 5 |
| T11P | 2 | 8 |
| T11R | 1 | 9 |
| S12G | 1 | 10 |
| S12P | 2 | 9 |
| H13F | 2 | 7 |
| P14M | 8 | - |
| R15A | >64 | - |
| I17M | 4 | 7 |
| R18A | 8 | 2 |
| Api137[a)] | 1 | 7 |

**Extended Data Table. 1 Cell-inhibitory activity of synthetic Dro variants**
The second column shows the result of the liquid culture MIC experiments in 25% cation-adjusted MHB medium. The third column shows the zone of no cell growth in the drop-diffusion test on the supplemented M9 minimal medium plate (Fig. 5b).

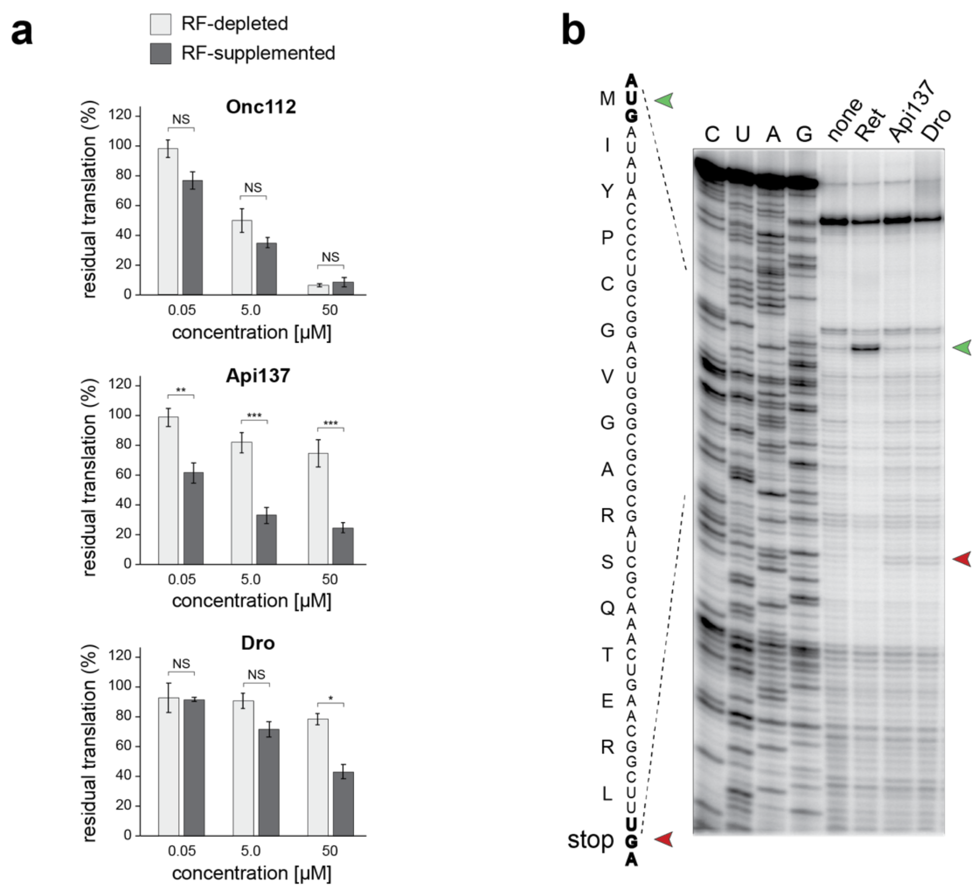


**Extended Data Fig. 1. Dro acts upon translation termination but causes weak translation arrest at UGA stop codons**

**a,** Inhibition of in vitro GFP translation by synthetic PrAMPs: class-I Oncocin112 (Onc112), class-II Api137, or non-glycosylated Dro (Dro). Bar graphs represent the normalized values of GFP fluorescence from reactions where RF1 was depleted (light grey) or supplemented (dark grey), setting the fluorescence value from reactions with or without RF1 in the absence of PrAMP as 100%. Error bars show deviation from the mean in three independent experiments. Significance levels indicated as NS, not significant; *, p-value <0.05; **, p-value < 0.01; ***, value < 0.001 (Tukey's range test). **b,** In vitro toeprinting analysis of the Api137 or Dro-mediated ribosome arrest at the UGA stop codon (red arrowhead) of the model *yrbA* ORF. The control reaction with no added PrAMPs is labeled as "none". The control antibiotic retapamulin (Ret) stalls ribosomes at the start codon (green arrowhead). Sequencing reactions are labeled as C, U, A, G.

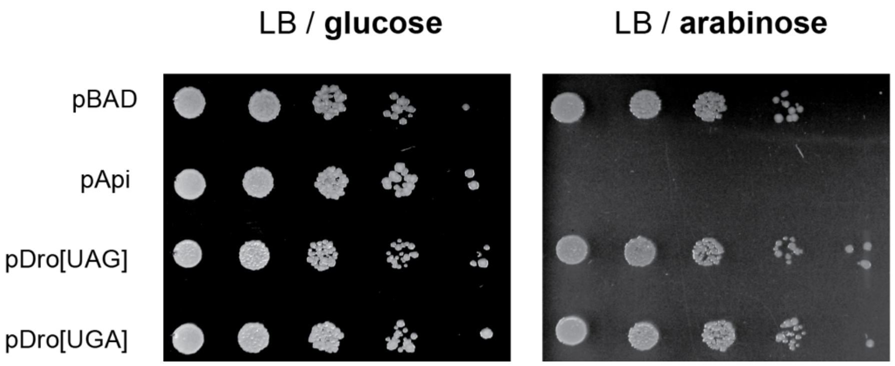


**Extended Data Fig. 2 Endogenous expression of Dro in cells grown in rich medium does not prevent cell growth**

Growth of *E. coli* BL21 cells transformed with pDro[UAG] or pDro[UGA] plasmids on agar lysogeny broth (LB) rich medium supplemented with glucose or L-arabinose. The toxic effect of endogenous expression of Api (in cells transformed with pApi) is shown for comparison. Cells transformed with empty pBAD vector were used as a negative control.

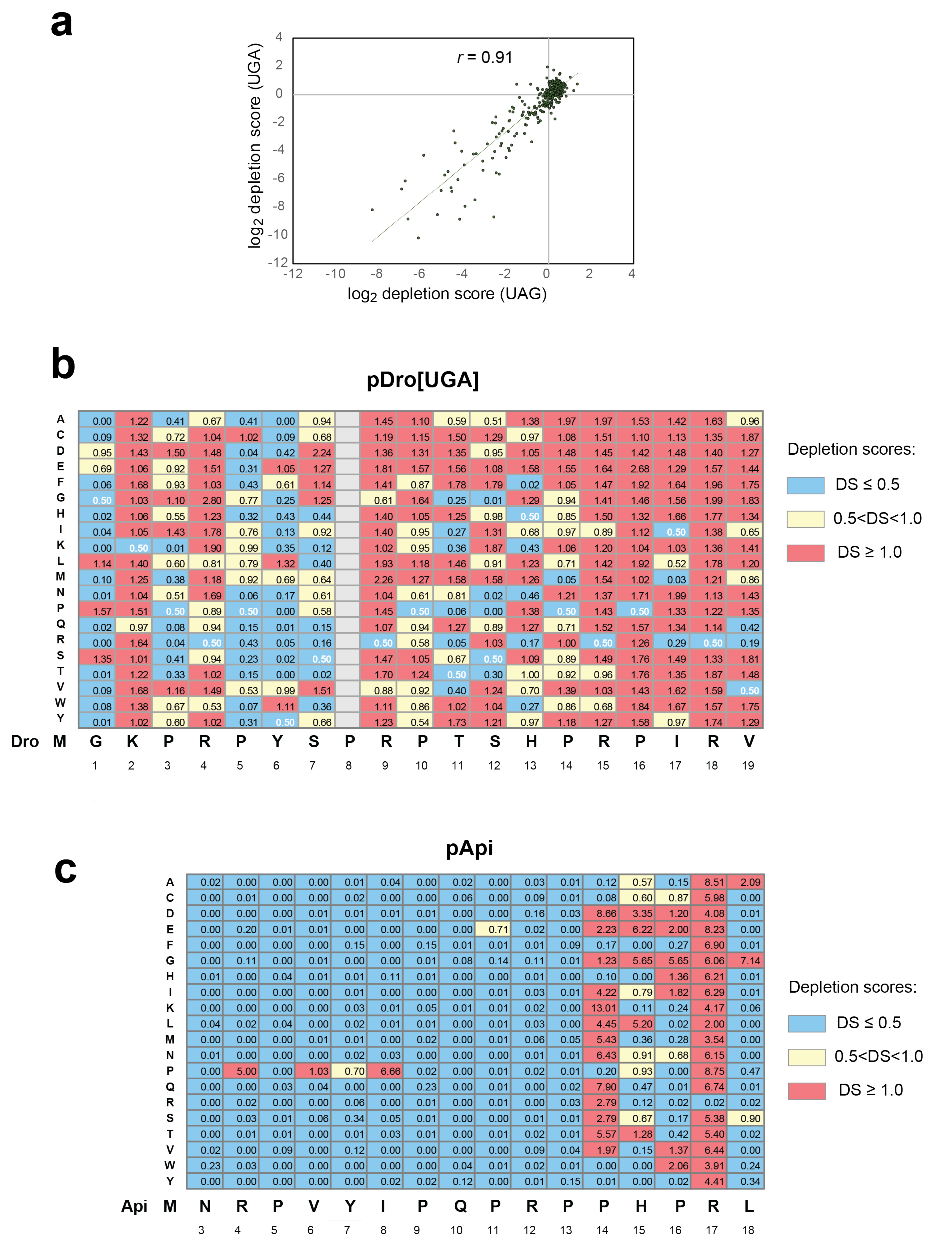


**Extended Data Fig. 3 Robustness of experimental data and comparisons of the effect of endogenously expressed Dro and Api variants on cell growth**

**a**, Correlation of the depletion scores of *E. coli* clones from the single amino acid Dro mutant libraries generated on the bases of pDro[UAG] or pDro[UGA] expression plasmids. Pearson correlation coefficient (*r*) is indicated. **b**, Depletion scores of library clones endogenously expressing single-amino acid Dro variants from the pDro[UGA] plasmids. Coloring is according to the toxic effects (blue - highly toxic, yellow - mildly toxic, salmon - not toxic). The analogous data for the pDro[UAG] library are shown in Fig. 4. **c,** Similarly calculated depletion scores for endogenously expressed Api mutants from the pApi plasmid using data from the reference [10].

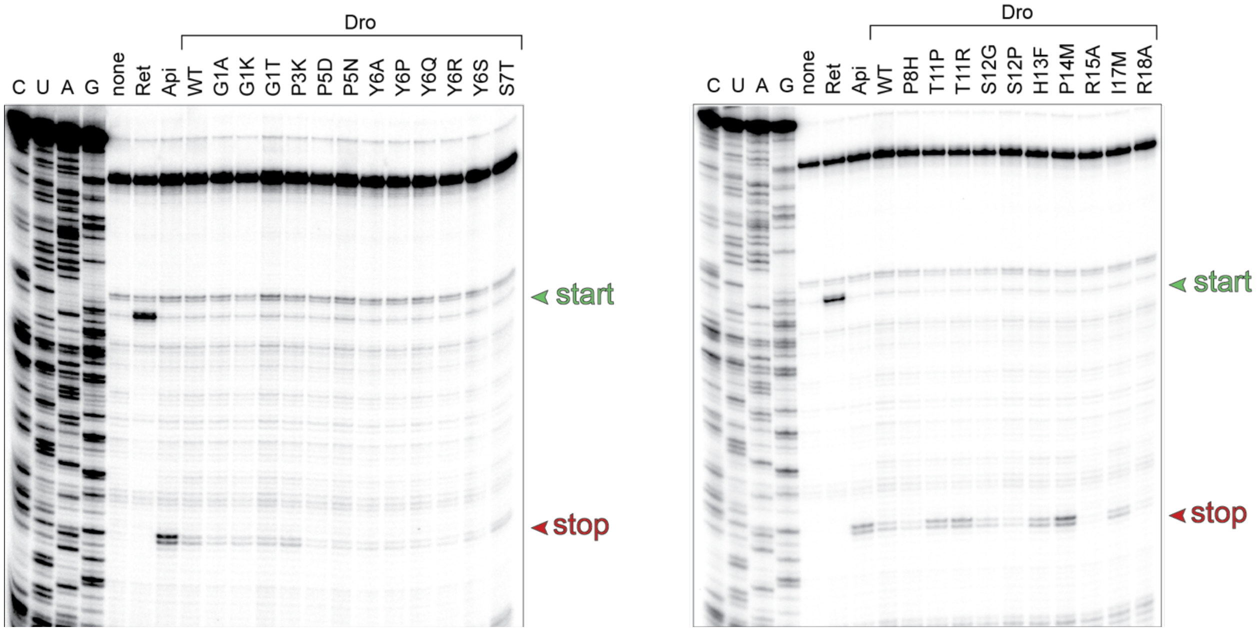


**Extended Data Fig. 4 Dro variants with single amino acid substitutions retain the ability to arrest ribosomes at stop codons**

Toeprinting analysis of the ribosome arrest of the UAG stop codon of the model *yrbA* ORF mediated by synthetic non-glycosylated Dro variants. Samples with no added synthetic peptide are labeled as 'none'. Arrest at stop codons caused by Api137 or at start codons by retapamulin (Ret) are shown as reference. Toeprint bands from ribosomes stalled at start or stop codons are marked by green and red arrowheads, respectively.

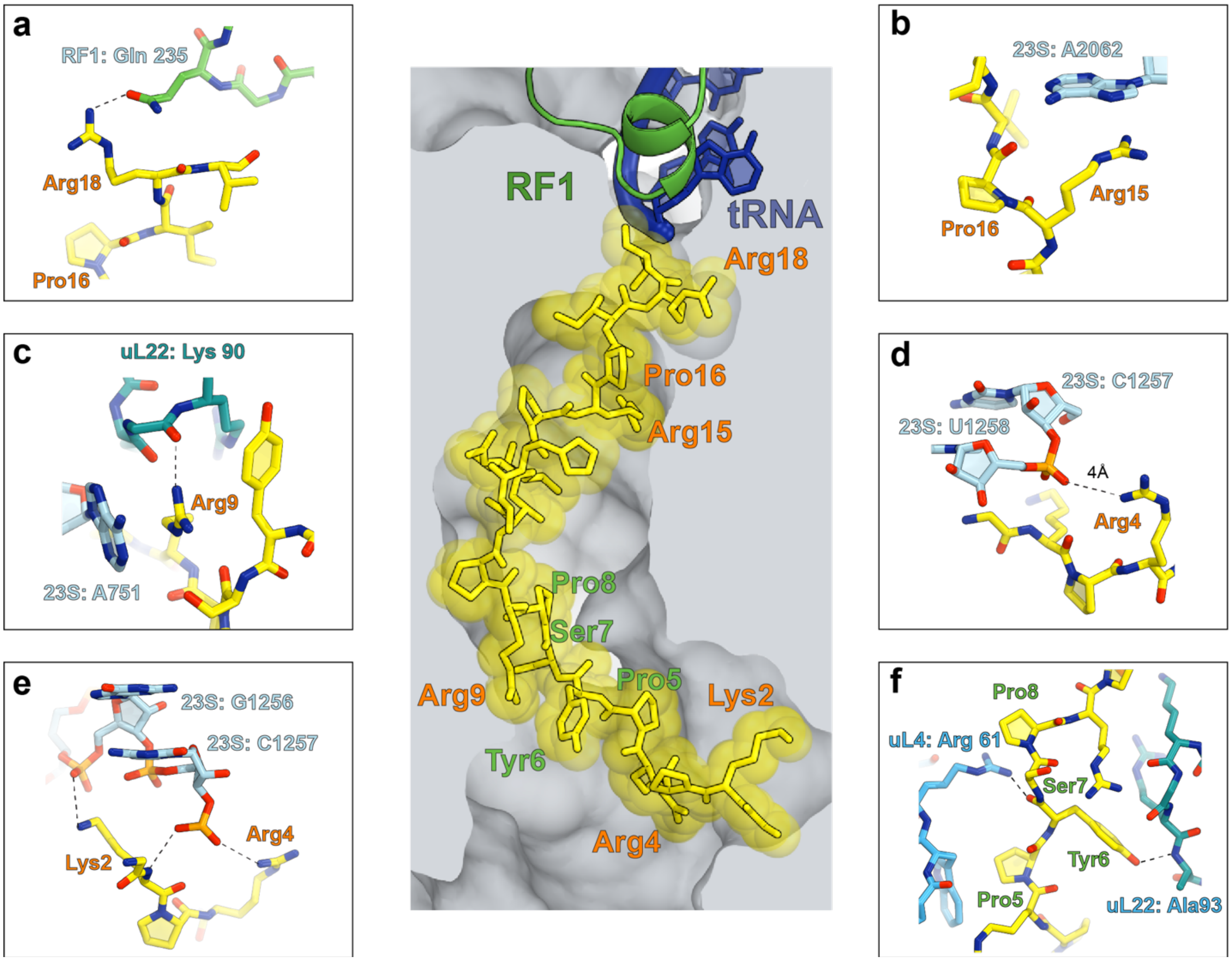


**Extended Data Fig. 5 Functionally critical contacts of Dro with the ribosome as revealed by mutational analysis.**
The central image depicts the placement of glycosylated Dro in the NPET of the *E. coli* ribosome [57]. Functionally critical Dro residues are indicated in salmon; residues that tolerate multiple substitutions are marked in green. **a-e**, Functionally critical contacts involving Dro residues: **a**, Arg18, **b**, Arg15 and Pro16, **c**, Arg9, **d**, Arg4, **e**, Lys2. Panel **f** shows the ribosomal contacts of the Pro5-Pro8 segment of Dro where most amino acids substitutions do not impair the PrAMP's activity.
